# Aging-Associated Decrease in the Histone Acetyltransferase KAT6B Causes Myeloid-Biased Hematopoietic Stem Cell Differentiation

**DOI:** 10.1101/679738

**Authors:** Eraj Shafiq Khokhar, Sneha Borikar, Elizabeth Eudy, Tim Stearns, Kira Young, Jennifer J. Trowbridge

**Affiliations:** Graduate School of Biomedical Science and Engineering, University of Maine, Orono, ME, 04469; The Jackson Laboratory, Bar Harbor, ME, 04609

## Abstract

Aged hematopoietic stem cells (HSCs) undergo biased lineage priming and differentiation toward production of myeloid cells. A comprehensive understanding of gene regulatory mechanisms causing HSC aging is needed to devise new strategies to sustainably improve immune function in aged individuals. Here, a focused shRNA screen of epigenetic factors reveals that the histone acetyltransferase *Kat6b* regulates myeloid cell production from hematopoietic progenitor cells. Within the stem and progenitor cell compartment, *Kat6b* is most highly expressed in long-term (LT)-HSCs and is significantly decreased with aging at the transcript and protein levels. Knockdown of *Kat6b* in young LT-HSCs causes skewed production of myeloid cells both *in vitro* and *in vivo*. Transcriptome analysis identifies enrichment of aging and macrophage-associated gene signatures alongside reduced expression of self-renewal and multilineage priming signatures. Together, our work identifies KAT6B as an epigenetic regulator of LT-HSC aging and a novel target to improve aged immune function.

## Introduction

Healthspan is defined as the period of time from birth until the organism remains free from chronic diseases (Kaeberlein, 2018). Aging involves a progressive decline in many cellular systems, resulting in a decrease in healthspan. Among these changes, reductions in immune system function have a significant contribution to shortening healthspan. As the immune system is responsible for detection and neutralization of pathogens and foreign particles (Parkin and Cohen, 2001), the elderly become more susceptible to infections, leading to more frequent and severe illness (Dorshkind, & Swain, 2009). With the global population of individuals aged 65 years and older expected to reach 1.6 billion by 2050 (Gasteiger et al., 2016), there is a pressing need to develop novel therapeutic strategies to ameliorate aging-associated declines in hematopoietic and immune function.

HSCs give rise to all mature blood cells. However, with age, cell-intrinsic changes within HSCs contribute to aging-associated hematopoietic decline (Rodrigues et al., 1997) such as increased HSC frequency, enhanced differentiation toward myeloid cells, and decreased ability to return to quiescence after activation (Verovskaya et al., 2019). Molecular features that contribute to these phenotypes include loss of cell polarity, impaired DNA damage repair, increased production of reactive oxygen species (ROS), and declines in mitochondrial function (Verovskaya et al., 2019). In addition, previous literature supports the occurrence of epigenetic drift in aged HSCs. This involves decreased expression of key epigenetic regulators, a global increase in DNA methylation (Beerman et al., 2013; Sun et al., 2014), and altered levels of histone H3 lysine 4 trimethylation (H3K4me3) and lysine 27 trimethylation (H3K27me3) in both aged murine and human HSCs (Adelman et al., 2019; Sun et al., 2014). Moreover, diminished levels and polarity of histone H4 lysine 16 acetylation (H4K16ac) is associated with loss of regenerative capacity and gain of myeloid lineage skewing in aged LT-HSCs (Florian et al., 2012). While these studies support involvement of epigenetic regulatory processes in HSC aging, there remains a lack of comprehensive knowledge of the extent to which epigenetic alterations cause aging-associated changes in HSC function. Here, we report a functional screen to uncover novel epigenetic regulators of aging-associated myeloid-biased differentiation from multipotent progenitor cells, identifying the lysine acetyltransferase *Kat6b*.

KAT6B (also known as MORF) belongs to the MYST family of histone acetyltransferases and is responsible for acetylation of the lysine 23 residue of histone H3 (H3K23ac) (Simó-Riudalbas et al., 2015). Other members of the MYST family, KAT6A (lysine acetyltransferase 6A; MOZ) and KAT8 (lysine acetyltransferase 8; MOF), have known functional roles in hematopoiesis. KAT6A, which catalyzes acetylation of lysine 9 (H3K9ac) and lysine 14 (H3K14ac) residues (Huang et al., 2016), is critical for the emergence and maintenance of hematopoietic stem cells (HSCs) (Katsumoto et al., 2006; Perez-Campo et al., 2009; Sheikh et al., 2016). KAT8, which catalyzes acetylation of the lysine 16 residue of histone H4 (H4K16ac), is critical for adult but not early fetal hematopoiesis (Valerio et al., 2017), supporting the concept that distinct histone acetyltransferases may be more critical at different ontological stages of hematopoiesis. Here, we investigate and demonstrate a novel role for KAT6B in lineage differentiation of LT-HSCs in the context of aging.

## Results

### An shRNA screen identifies *Kat6b* as a novel regulator of myeloid differentiation from lymphoid-primed multipotent progenitor cells

To identify epigenetic regulators that have a functional role in aging-associated myeloid lineage-biased differentiation of hematopoietic stem and progenitor cells (HSPCs), we conducted an *in vitro* shRNA screen. Using gene expression commons (GEXC) (Seita et al., 2012), we identified 2,766 differentially expressed genes between granulocyte macrophage progenitors (GMPs) and common lymphoid progenitors (CLPs) (Figure 1A), which are committed progenitors for the myeloid and lymphoid lineages, respectively (Motonari, 2013). Among these 2,766 differentially expressed genes, gene ontology (GO) enrichment analysis of Reactome pathways (Carbon et al., 2019; Mi et al., 2017; The Gene Ontology Consortium et al., 2000) revealed significant enrichment of chromatin modifying enzymes (P = 0.000158, FDR = 0.00111). The 40 enriched genes encoding chromatin modifying enzymes were further subset to 30 genes based on overlap with the GO annotation “regulation of gene expression” (GO:0010468) (Supplemental Figure 1A). Lastly, this gene list was filtered to include those with commercially available shRNA constructs with verified knockdown in murine cell lines, resulting in 16 genes (Supplemental Table 1). To begin functional screening, shRNA expression plasmids for six of these 16 genes were obtained. For negative and positive controls, we used a scrambled shRNA-expressing non-targeting control (NTC) vector and a shRNA vector targeting CREB-binding protein (*Crebbp*), respectively. Conditional knockout of *Crebbp* is known to cause loss of HSPCs and result in myeloid-biased hematopoiesis (Chan et al., 2011). In addition, shRNA constructs were obtained for eight genes hypothesized to regulate lineage differentiation using a candidate gene approach (Supplemental Table 2). After cloning, we validated reduced target gene expression from each of these shRNA constructs in murine 3T3 cell lines (Supplemental Figure 1B).

**Figure 1.**
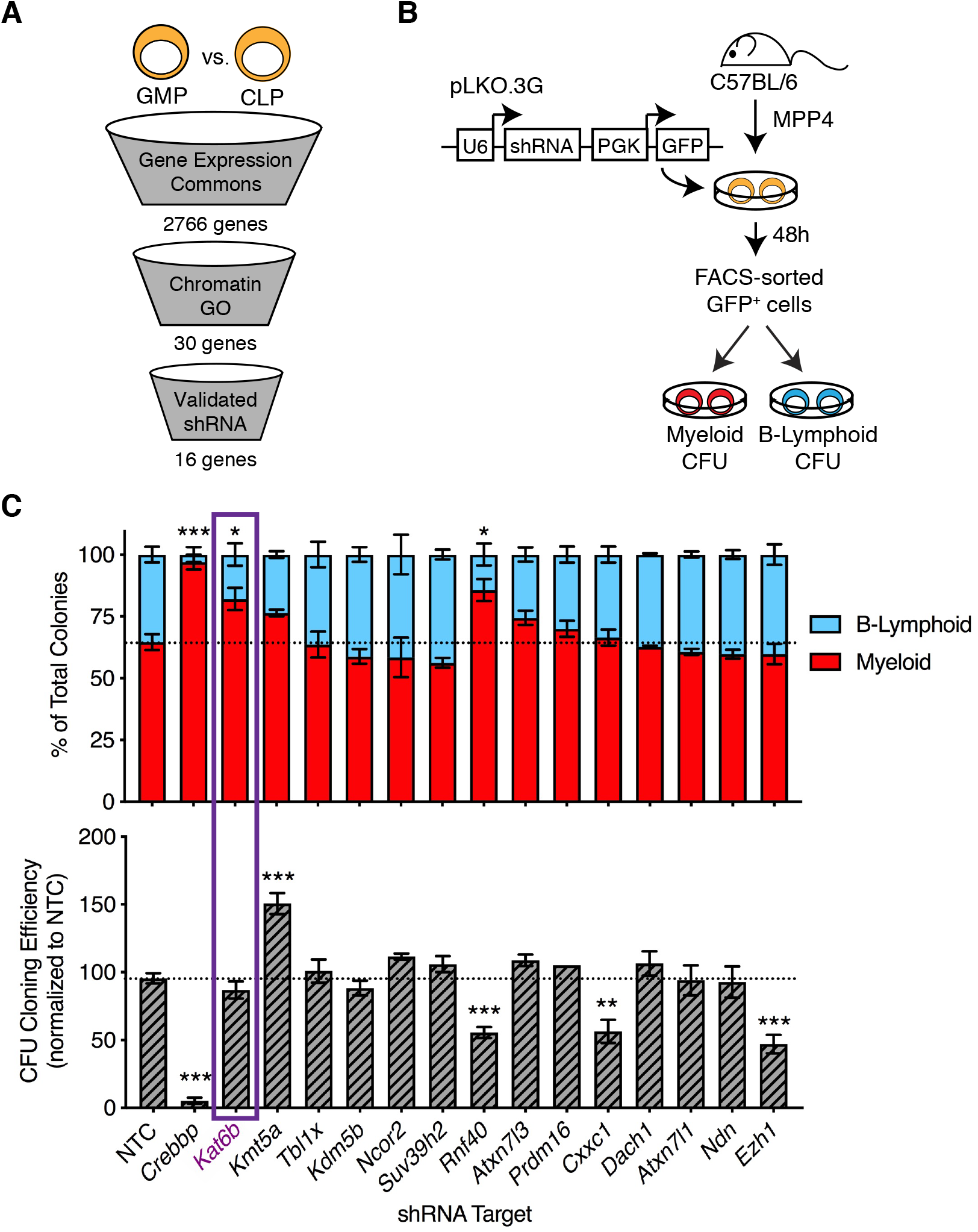
Functional shRNA screen for epigenetic regulators of myeloid versus B-lymphoid differentiation identifies *Kat6b*. (A) Schematic of candidate selection criteria to identify chromatin regulatory genes involved in myeloid versus B-lymphoid differentiation of hematopoietic stem and progenitor cells. GMP; granulocyte-macrophage progenitors, CLP; common lymphoid progenitors. (B) Schematic of experimental design to test epigenetic regulatory gene candidates using shRNA-mediated knockdown in lymphoid-primed multipotent progenitor cells (MPP4) and colony-forming unit (CFU) assays. (C) (Top panel) Frequency of myeloid and B-lymphoid colonies out of total colonies and (Bottom panel) CFU cloning efficiency calculated as the total number of myeloid and B-lymphoid colonies following shRNA knockdown of the indicated target genes divided by the total number of myeloid and B-lymphoid colonies in NTC. NTC; non-targeting control. Bars represent mean ± SEM of *n* = 2 biological replicates. \*\**P* < 0.01, \*\*\**P* < 0.001 by Holm-Sidak’s multiple comparisons test.

To perform the *in vitro* shRNA screen, we utilized lymphoid-primed multipotent progenitor cells (MPP4) as our starting cell population. MPP4 cells were selected as they have both lymphoid and myeloid differentiation potential (Pietras et al., 2015) and, in contrast to LT-HSCs, have efficient clonal *in vitro* differentiation capacity giving rise to both lymphoid and myeloid cells (Young et al., 2016). MPP4 cells were isolated by fluorescence-activated cell sorting (FACS) from young adult (8-10 weeks old) mice, transduced with lentiviral particles containing individual shRNA expression plasmids, and cultured for two days with growth factors that we previously identified as supporting both lymphoid and myeloid differentiation from this population (Young et al., 2016) (Figure 1B). After two days, GFP-expressing cells were isolated by FACS and plated into parallel myeloid and B-lymphoid colony-forming unit (CFU) differentiation assays. To identify genes responsible for myeloid versus B-lymphoid differentiation, we sought genes whose knockdown produced a significant change in the proportion of myeloid relative to B-lymphoid colonies while maintaining overall cloning efficiency. Relative to NTC, we found that knockdown of our positive control *Crebbp* resulted in a near-complete loss of CFU capacity and the residual colonies that formed were predominantly myeloid (Figure 1C), consistent with the expected phenotype of *Crebbp* loss (Chan et al., 2011). In two out of the 14 shRNA constructs evaluated, targeting *Kat6b* and *Rnf40* (ring finger protein 40), we observed a significant increase in the proportion of myeloid relative to B-lymphoid colonies. Of these, only knockdown of *Kat6b* was found not to alter overall cloning efficiency and thus was pursued as a candidate epigenetic regulator of aging-associated myeloid lineage bias.

### KAT6B decreases at the transcript and protein level in aged LT-HSCs

To gain insight into the functional role of *Kat6b* in aging-associated myeloid lineage-biased differentiation of HSPCs, we sought to determine where *Kat6b* is most abundantly expressed within the HSPC compartment and how this expression may be altered in aging. We mined publicly available RNA-seq data (Lara-Astiaso et al., 2014). We observed the highest transcript level of *Kat6b* in LT-HSCs relative to other stem and progenitor cell populations (Figure 2A). To determine whether *Kat6b* expression is altered in the context of aging, we isolated LT-HSCs and MPP4 cells by FACS from young (2-4 month) and aged (20-23 month) mice. By real-time PCR, we observed that the *Kat6b* transcript decreases 2.8-fold with age in LT-HSCs (Figure 2B) but does not decrease with age in MPP4 cells (Supplemental Figure 2A). Previous work comparing transcriptional changes between young and aged mouse LT-HSCs found a 1.2-fold decrease in *Kat6b* expression in aged LT-HSCs (FDR = 0.0413) (Sun et al., 2014) and a recent study comparing HSCs isolated from young (18-30 year-old) and aged (65-75 year-old) humans (Adelman et al., 2019) identified that *KAT6B* transcript decreases 1.2-fold in aging (Padj = 0.0397) (Supplemental Figure 2B), supporting our finding that *Kat6b* levels decrease with age in HSCs. To analyze KAT6B at the protein level, we immuno-stained LT-HSCs isolated by FACS from young and aged mice with an antibody against KAT6B and the nuclear stain DAPI (Figure 2C). We observed that the mean fluorescence intensity (MFI) of KAT6B in LT-HSCs isolated from aged mice is significantly lower than in young mice (Figure 2D). Together, our results show that KAT6B is significantly decreased at both the transcript and protein levels in aged LT-HSCs.

**Figure 2.**
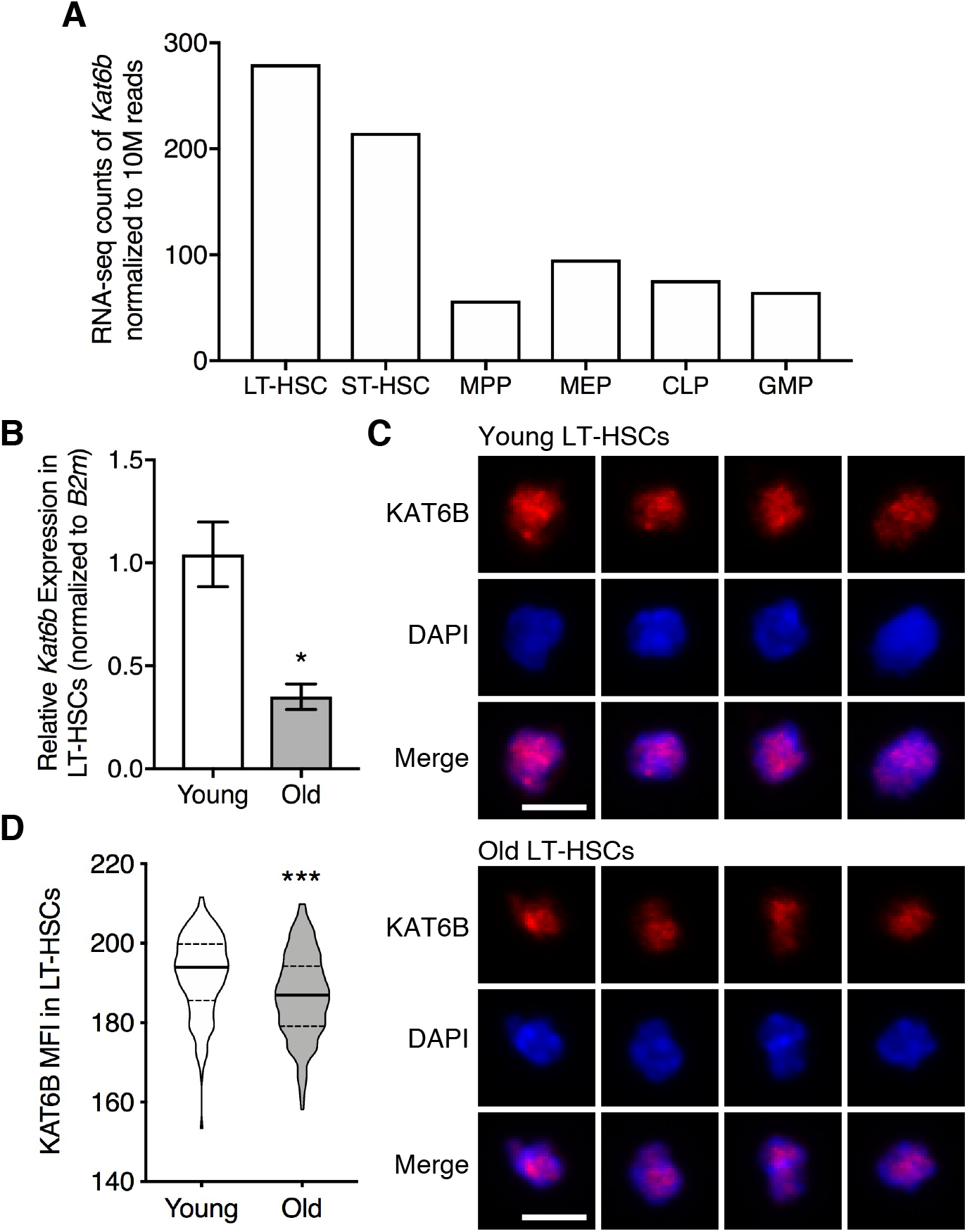
KAT6B is decreased in aged LT-HSCs. (A) Relative expression of *Kat6b* in hematopoietic stem and progenitor cell populations (data from Lara-Astiaso et al., 2014). (B) Relative expression of *Kat6b* in LT-HSCs isolated from young (2-4 month) and aged (20-23 month) mice. Bars represent mean ± SEM of *n* ≥ 3 biological replicates. \**P* < 0.05 by unpaired *t* test. (C) Representative immunofluorescence images of KAT6B and DAPI in LT-HSCs isolated from young and aged mice. Scale bar equals 5 um. (D) Violin plots of mean fluorescence intensity (MFI) of KAT6B in LT-HSCs isolated from young and aged mice. Solid lines indicate median and dotted lines indicate quartiles. Data points include *n* = 29-45 individual cells sampled from *n* = 4 biological replicate animals. \*\*\**P* < 0.001 by unpaired *t* test.

### Knockdown of *Kat6b* in LT-HSCs causes myeloid-biased *in vitro* differentiation in CFU assays

To evaluate the functional consequence of reduced expression of *Kat6b* as observed in aged LT-HSCs, we utilized a shRNA knockdown approach. LT-HSCs isolated from young mice were transduced with lentiviral particles containing NTC or one of two *Kat6b* shRNA expression plasmids and cultured for two days with growth factors supporting LT-HSC maintenance (Figure 3A) (Holmfeldt et al., 2016). After two days, GFP^+^ cells were isolated by FACS and evaluated for *in vitro* myelo-erythroid differentiation using CFU assays. From the resultant colonies, we determined that *Kat6b* transcript was reduced by 5.7-fold and 1.4-fold using *Kat6b* shRNA1 (sh1) and *Kat6b* shRNA2 (sh2), respectively (Figure 3B). The total number of colonies was not significantly altered in sh1 or sh2 compared to NTC (Figure 3C). However, differences were observed with respect to colony composition, determined based upon cellular morphology within each colony to distinguish macrophage-only (CFU-M), granulocyte-macrophage (CFU-GM) and granulocyte-erythrocyte-macrophage-megakaryocyte (CFU-GEMM) colonies. Upon knockdown of *Kat6b*, we observed a significant increase in the number of CFU-GM colonies and a significant decrease in the number of CFU-GEMM colonies (Figure 3D), consistently with both shRNA constructs.

**Figure 3.**
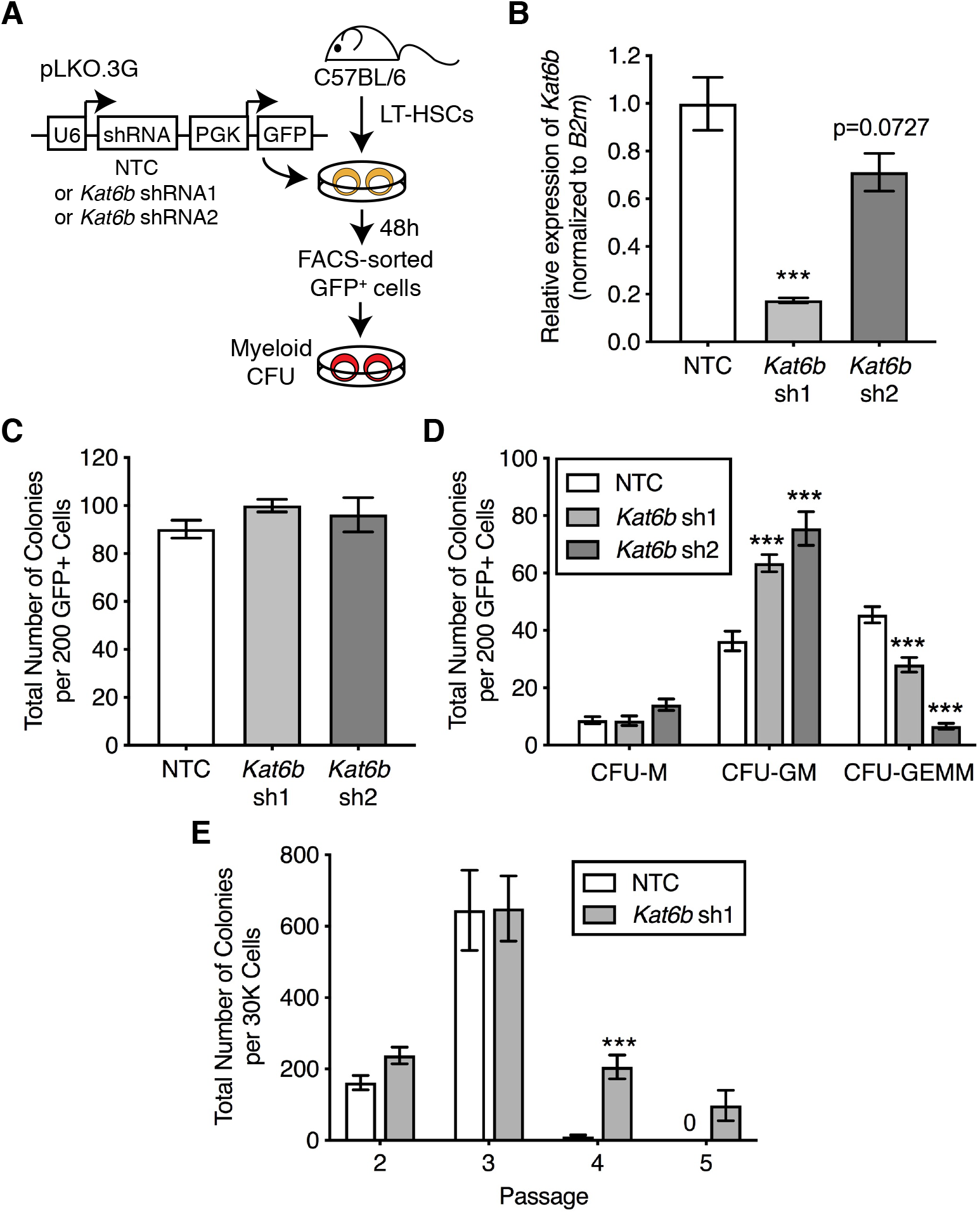
*Kat6b* knockdown alters myeloid differentiation of LT-HSCs *in vitro*. (A) Schematic of experimental design to knockdown *Kat6b* in LT-HSCs and assess differentiation in the myeloid CFU assay. (B) Relative expression of *Kat6b* in colonies following shRNA-mediated knockdown of *Kat6b* using two independent hairpins (sh1 or sh2) or NTC. (C) Total number of colonies produced and (D) colony subtype distribution from 200 GFP^+^ cells posttransduction of LT-HSCs. CFU-M; macrophage, CFU-GM; granulocyte-macrophage, CFU-GEMM; granulocyte-erythrocyte-macrophage-megakaryocyte. (E) Total number of colonies produced upon passage of 30K cells harvested from the primary CFU assay. In all graphs, bars represent mean ± SEM of *n* ≥ 3 biological replicates. \*\*\**P* < 0.001 by Holm-Sidak’s multiple comparisons test.

In addition, we investigated the effect of *Kat6b* knockdown on colony replating capacity, an *in vitro* surrogate of self-renewal. While NTC colonies did not replate past passage three, we observed that *Kat6b* knockdown colonies replated to passage five, representing a significant increase in CFU replating capacity (Figure 3E). Together, our results demonstrate that knockdown of *Kat6b* results in myeloid-biased *in vitro* differentiation from LT-HSCs and increased serial replating capacity of the resultant myeloid progenitor cells.

### Knockdown of *Kat6b* in LT-HSCs causes myeloid-biased differentiation *in vivo*

To evaluate the functional consequence of reduced levels of *Kat6b* in LT-HSCs *in vivo*, we transduced LT-HSCs with *Kat6b* sh1 or NTC and transplanted GFP^+^ cells into lethally irradiated B6.CD45.1 recipient mice (Figure 4A). In total, 15 recipient mice were transplanted with NTC-transduced cells and 16 recipient mice were transplanted with *Kat6b* sh1-transduced cells. From these, 7/15 (46%) and 8/16 (50%) were found to have multilineage engraftment above a threshold of 0.1% donor-derived peripheral blood cells at one month post-transplant. At this time point, donor-derived engraftment (% CD45.2^+^ GFP^+^) was not significantly different between NTC and *Kat6b* sh1 (Figure 4B). However, mice transplanted with *Kat6b* knockdown LT-HSCs had a significant increase in the proportion of donor-derived myeloid cells in the peripheral blood as compared to NTC (Figure 4C). In addition, there was a significant decrease in donor-derived erythroid cells in the peripheral blood of mice transplanted with *Kat6b* knockdown LT-HSCs compared to NTC (Figure 4D). A trend toward decreased frequency of donor-derived B and T lymphocytes in *Kat6b* knockdown compared to NTC did not reach statistical significance (*P* = 0.1264 and *P* = 0.1735, respectively) (Supplemental Figure 3A).

**Figure 4.**
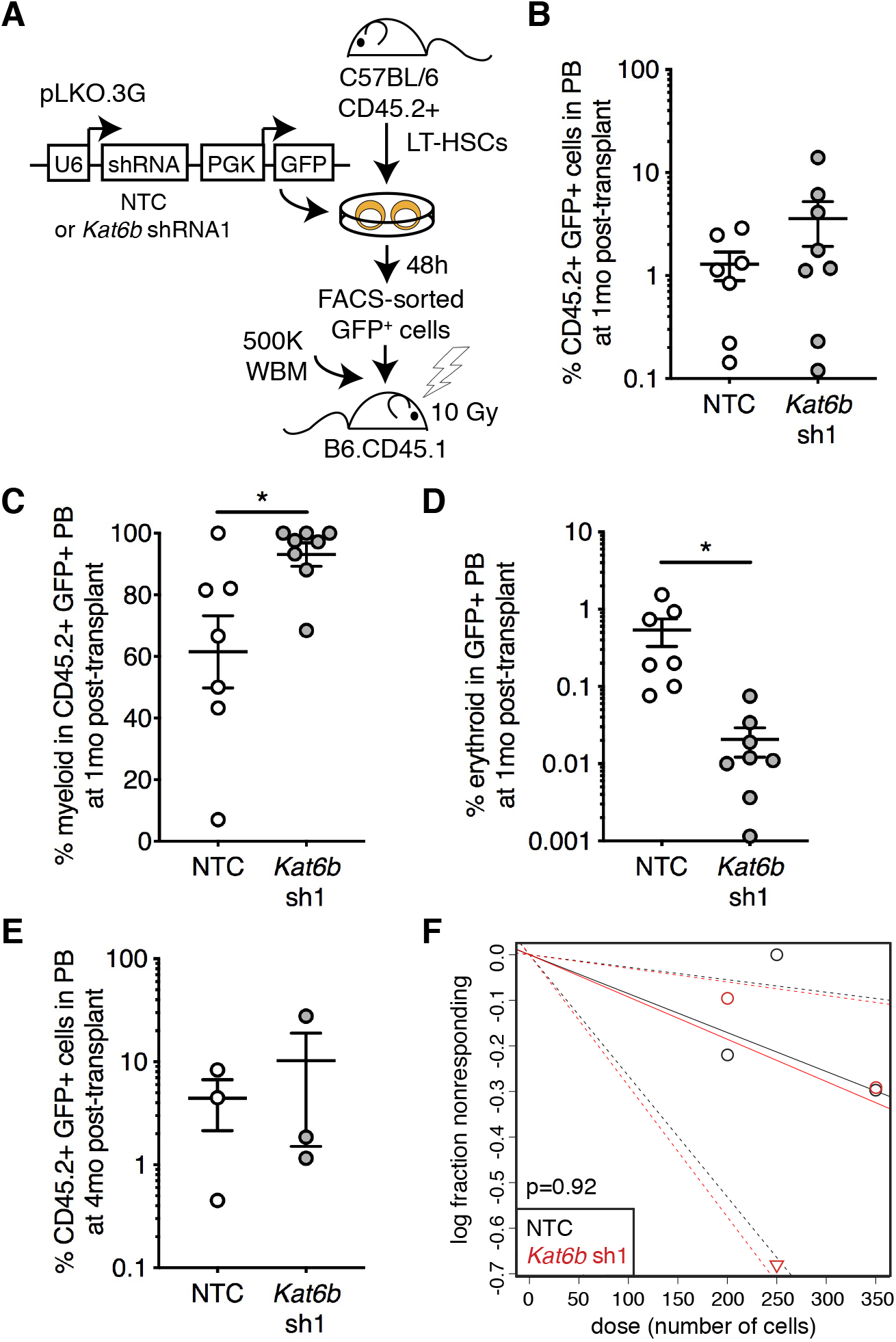
*Kat6b* knockdown alters myeloid and erythroid differentiation of LT-HSCs *in vivo*. (A) Schematic of experimental design to knockdown *Kat6b* in LT-HSCs and assess hematopoietic reconstitution in lethally irradiated recipient mice compared to NTC-transduced LT-HSCs. (B) Frequency of donor-derived cells (CD45.2^+^ GFP^+^) in the peripheral blood (PB) of recipient mice, (C) myeloid cells (CD11b+) within donor-derived PB cells (CD45.2^+^ GFP^+^), and (D) erythroid cells (Ter119^+^) within donor-derived PB cells (GFP^+^) at 1 month (1mo) posttransplant. Each dot represents one recipient mouse. Lines represent mean ± SEM of *n* ≥ 7 biological replicates. \**P* < 0.05 by unpaired *t* test (E) Frequency of donor-derived cells in the peripheral blood (PB) of recipient mice at 4mo post-transplant. Each dot represents one recipient mouse. Lines represent mean ± SEM of *n* = 3 biological replicates. (F) Limiting dilution analysis of repopulating cell frequency from NTC or *Kat6b*-transduced LT-HSCs at 4mo post-transplant.

At four months post-transplant, 3/15 (20%) and 3/16 (18.8%) of recipients were found to have sustained multilineage engraftment above a threshold of 0.1% donor-derived peripheral blood cells (Figure 4E, Supplemental Figure 3B). These data were utilized to calculate repopulating cell frequency by limiting dilution analysis (Hu and Smyth, 2009). In the NTC group, the repopulating cell frequency was calculated to be 1/1170 (1/3644 to 1/376; 95% Confidence Interval (CI)), similar to the repopulating cell frequency in *Kat6b* knockdown (1/6720, 1/1683 to 1/422; 95% CI) (Figure 4F). Together, these results show that knockdown of *Kat6b* causes myeloid-biased differentiation from LT-HSCs *in vivo* without altering repopulation capacity.

### Knockdown of *Kat6b* in LT-HSCs Decreases Multilineage Priming and Promotes Expression of Inflammation-Associated Gene Signatures

To investigate the molecular mechanisms underlying myeloid-biased differentiation after *Kat6b* knockdown, we transduced LT-HSCs with NTC or *Kat6b* sh1 and performed RNA-seq on sorted GFP^+^ cells. Unsupervised clustering separated NTC and *Kat6b* knockdown samples (Figure 5A). 252 significantly differentially expressed genes (*FDR* < 0.05) were identified, out of which 127 genes were upregulated and 125 genes were downregulated in *Kat6b* knockdown compared to NTC (Figure 5B). *Kat6b* itself was found to be downregulated by 1.2-fold in *Kat6b* knockdown compared to NTC samples. In addition, we observed downregulation of *Apoe* (apolipoprotein E), loss of which impairs B cell development and HSC function (Gasparetto et al., 2012) and *Aldh3a1* (aldehyde dehydrogenase family 3, subfamily A1), loss of which has been demonstrated to cause monocytosis and neutrophilia in mice (Murphy et al., 2011).

**Figure 5.**
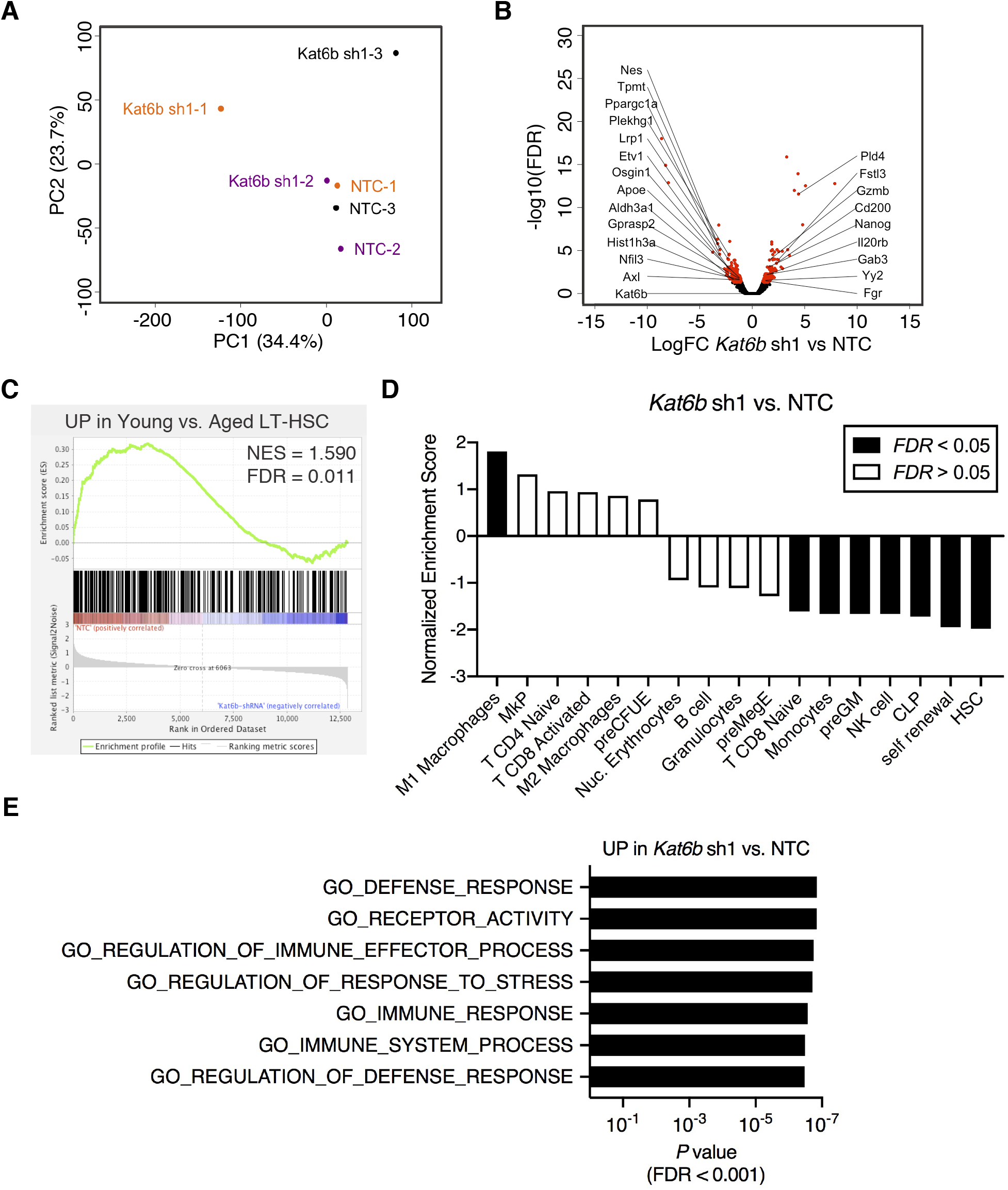
*Kat6b* knockdown alters gene expression programs critical for multilineage differentiation. (A) PCA plot showing unsupervised clustering of gene expression profiles from *Kat6b* sh1 (*n* = 3) and NTC (*n* = 3). Each color represents a set of biological replicate samples. (B) Volcano plot showing log fold changes of genes against −log10 of *FDR*. Points in red highlight genes with *FDR* < 0.05. (C) Gene set enrichment analysis (GSEA) of genes upregulated in young versus aged LT-HSCs (Sun et al., 2014). Red denotes NTC and blue denotes *Kat6b* sh1. (D) Normalized enrichment score from GSEA analysis of the indicated datasets in *Kat6b* sh1 versus NTC. Black bars indicate *FDR* < 0.05, white bars indicate *FDR* > 0.05. (E) Top gene ontology (GO) terms enriched in genes found to be significantly upregulated in *Kat6b* sh1 versus NTC (fold change > 2 and *P* < 0.05).

To test the hypothesis that *Kat6b* knockdown alters expression of gene programs associated with aging and differentiation of LT-HSCs, we performed gene set enrichment analysis (GSEA) (Daly et al., 2003; Subramanian et al., 2005). Comparing our RNA-seq data to LT-HSC aging gene signatures (Sun et al., 2014) revealed that genes more highly expressed in young versus aged LT-HSCs were significantly enriched in NTC versus *Kat6b* knockdown (Figure 5C). We then compared our dataset to previously defined gene signatures representing HSCs (Chambers et al., 2007), the self-renewal program (Krivtsov et al., 2006), hematopoietic progenitor cell populations (lymphoid (CLP), granulocyte-macrophage (preGM) and erythroid-megakaryocyte (preMegE, preCFU-E, MkP)) (Sanjuan-Pla et al., 2013), and mature hematopoietic cell populations (M1 and M2 macrophages, monocytes, granulocytes, erythrocytes, CD4+ naïve T cells, CD8+ naïve and activated T cells, B cells and NK cells) (Chambers et al., 2007; Engler et al., 2012; Mantovani et al., 2002; Martinez et al., 2006) (Supplemental Table 3). This analysis revealed that *Kat6b* knockdown LT-HSCs had a significant enrichment of M1 macrophage signatures while NTC LT-HSCs were enriched in HSC/self-renewal, preGM, monocyte, CLP, NK and CD8+ naïve T cell signatures (Figure 5D). To further interrogate mechanisms underlying the observed myeloid differentiation bias of *Kat6b* knockdown LT-HSCs, unbiased GO enrichment analysis was utilized. This analysis revealed significant upregulation of signatures associated with defense response, immune processes and immune response in *Kat6b* knockdown LT-HSCs (Figure 5E) and downregulation of signatures associated with response to external stimulus and homeostasis (Supplemental Figure 4). Together, these data suggest that decreased expression of *Kat6b* in LT-HSCs impairs multilineage differentiation, and permits a transcriptional program promoting differentiation toward pro-inflammatory-type macrophages and that is associated with aging.

As KAT6B is known to catalyze H3K23 acetylation, we hypothesized that this modification would be decreased in LT-HSCs with aging. To test this hypothesis, we isolated LT-HSCs from young and aged mice by FACS and immunostained with an antibody against H3K23ac and DAPI (Supplemental Figure 5A). We observed a trend toward decrease in mean fluorescence intensity of H3K23ac in LT-HSCs isolated from aged mice (*P* = 0.1729) (Supplemental Figure 5B). These results suggest that H3K23ac may be modestly reduced with age in LT-HSCs, consistent with reduction in KAT6B at the transcript and protein levels.

## Discussion

In this study, by employing a shRNA-mediated screen of epigenetic regulators, we have discovered a novel role for *Kat6b* in the context of LT-HSC differentiation with relevance to aging. We have found that KAT6B decreases in aged HSCs at the transcript and protein levels. Knockdown of *Kat6b* in young LT-HSCs resulted in an increase in the proportion of myeloid cells and decrease in the proportion of erythroid cells *in vitro* and *in vivo*, demonstrating myeloid lineage-biased differentiation. Transcriptome data revealed that knockdown of *Kat6b* resulted in loss of HSC-associated expression signatures as well as multilineage priming, while gaining a M1 pro-inflammatory macrophage expression signature. Together, our results support that *Kat6b* functions as a regulator of HSC self-renewal and multilineage differentiation and decrease in *Kat6b*, as observed in aging, promotes myeloid lineage-biased HSC differentiation.

Our work builds upon literature demonstrating the importance of the MYST family of acetyltransferases for LT-HSC function. KAT6A, a paralogue of KAT6B with structural similarity (Simpson et al., 2012), also has important functions in regulation of hematopoietic stem and progenitor cells. However, our results support non-redundant roles for KAT6A and KAT6B in hematopoiesis. *In vitro, Kat6a*-deficient bone marrow has reduced total colony-forming units with no change in the ratio of individual colony types (Sheikh et al., 2016), whereas we observed that *Kat6b* knockdown results in no change in total colony numbers but a specific increase in the proportion of myeloid-only colonies. Furthermore, conditional deletion of *Kat6a* resulted in reduced total number of B cells and no change in total number of neutrophils and RBCs (Sheikh et al., 2016), whereas our *in vivo* experiments show that *Kat6b* knockdown resulted in no significant change in B cell frequency, increased frequency of myeloid cells and decreased frequency of RBCs. Thus, we propose that KAT6A and KAT6B have overlapping but distinct roles in proper multilineage differentiation from LT-HSCs.

In the context of our experiments, LT-HSCs were cultured under *ex vivo* conditions which have been reported to promote HSC self-renewal (Holmfeldt et al., 2016). However, this requirement for *ex vivo* culture for lentiviral transduction is also a caveat in the interpretation of our results. It is possible that some or all of the LT-HSCs seeded into *ex vivo* culture differentiate to progenitors during the 48h transduction culture period. Thus, the *Kat6b* knockdown phenotype we observe may be manifest in either HSCs or their myeloid progenitor progeny.

As a proof of principle that aged HSCs can be rejuvenated by therapeutic interventions that target epigenetic regulatory processes, it has been shown that inhibition of CDC42 by CASIN restored the polarity of H4K16ac, Cdc42 and Tubulin in aged HSCs (Florian et al., 2012). These CASIN-treated aged HSCs had re-balanced B-lymphoid and myeloid differentiation potential *in vivo* and HSC frequency resembling young animals (Florian et al., 2012). In addition, reprogramming of aged HSCs to induced pluripotent stem cells (iPSCs) has been shown to rejuvenate their *in vivo* engraftment and T cell differentiation potential (Wahlestedt et al., 2013). Our work suggests that therapeutically increasing levels of KAT6B in aged HSCs may also rejuvenate aspects of altered functionality, particularly with respect to lineage-balanced differentiation. A recent report by Adelman et al. demonstrated a reduction in active enhancer-associated chromatin modifications at a *KAT6B*-proximal enhancer region in aged versus young human HSCs (Adelman et al., 2019), suggesting that therapeutic approaches to increase enhancer activity may be a viable strategy to boost *Kat6b* expression in aged HSCs. Further studies will be required to test whether restoring expression of *Kat6b* in aged HSCs to levels observed in young HSCs is sufficient to restore balanced lineage differentiation.

## Experimental Procedures

### Experimental Animals

Young C57BL/6J (2-4 months old), aged C57BL/6J (20-24 months old) and B6.SJL-*Ptprc*^a^*Pepc*^b^/BoyJ (B6.CD45.1) (2-4 months old) were obtained from, and aged within, The Jackson Laboratory. All mice used in this study were females. All experiments were approved by The Jackson’s Laboratory Institutional Animal Care and Use Committee (IACUC).

### Plasmids

shRNA expression plasmids (in pLKO.1 or pLKO1.5) were obtained from Sigma (Supplemental Table 4). pLKO.3G (Addgene) was a gift from Christophe Benoist & Diane Mathis (Addgene plasmid # 14748; http://n2t.net/addgene:14748; RRID:Addgene_14748). The GFP cassette from pLKO.3G was sub-cloned into each shRNA plasmid. Colony PCR with eGFP primers (Supplemental Table 5) was performed to identify clones containing the GFP insert.

### Lentiviral Supernatant

shRNA expression plasmids, *RC-CMV Rev1b*, HDM-Hpgm2 (*gag-pol*), HPM-tat1b, HDM-*(VSV-G)* (at a mass ratio of 5:2:2:2:1) were transfected into 5 x 10^6^ HEK-293T cells, seeded 24 hours before transfection. Calcium phosphate transfection kit (Takara) was used for transfection. Growth media was replaced 24 hours after transfection. For viral supernatant collection media collected 48 hours after transfection, centrifuged at 1250 rpm 5 min 4°C and aliquots of supernatant were stored at −80°C. To titer lentiviral supernatant, 2 x 10^5^ NIH/3T3 cells were seeded in a 6-well plate 24 hours before supernatant was thawed and added at 1:2, 1:10, 1:50, 1: 250 and 1:1250 diluted in DMEM+10%FBS with 5 ug/ml polybrene. Plate was spun at 2500 rpm for 60 min at room temperature followed by incubation at 37°C and 5% CO_2_ for 48 hours with media change after 24 hours. After this 48h culture period, cells were harvested and run on a LSRII (BD Biosciences) to assess frequency of GFP^+^ cells. These frequencies were used to calculate the titer of each preparation of lentiviral supernatant. In addition, RNA was isolated from transduced NIH/3T3 cells to assess shRNA knockdown efficiency by real-time PCR.

### Primary Cell Isolation

LT-HSCs and MPP4 cells were isolated as described previously (Young et al., 2016). Femurs, tibiae, iliac crests from each mouse were pooled, crushed and filtered to prepare single-cell suspensions of bone marrow (BM). Ficoll-Paque (GE Healthcare Lifesciences) density centrifugation was used to isolate BM mononuclear cells (MNCs). These cells were stained with fluorochrome-conjugated antibodies from BioLegend, eBiosciences, BD Biosciences: c-Kit (clone 2B8), CD48 (clone HM48-1), CD150 (clone TC15-12F12.2), Sca-1 (clone 108129), FLT3 (clone A2F10) mature lineage (Lin) marker mix (B220 (clone RA3-6B2), CD11b (clone M1/70), CD4 (clone RM4-5), CD8a (clone 53-67), Ter-119 (clone TER-119), Gr-1 (clone RB6-8C5)) and viability stain propidium iodide (PI). Cells were sorted on a FACSAria (BD Biosciences) with these surface marker profiles: LT-HSC (Lin^−^ Sca^+^ c-Kit^+^ Flt3^−^ CD150^+^ CD48^−^) and MPP4 cells (Lin^−^ Sca^+^ c-Kit^+^ Flt3^+^).

### Transduction of LT-HSCs and MPP4 Cells

LT-HSCs were resuspended in SFEMII (Stemcell Technologies) supplemented with growth factors described previously (Holmfeldt et al., 2016); Stem cell factor (SCF; 10 ng/ml), thrombopoietin (TPO; 20 ng/ml), insulin-like growth factor 2 (IGF2; 20 ng/ml) and fibroblast growth factor (FGF; 10 ng/ml) (BioLegend or StemCell Technologies) along with 5ug/ml polybrene (Sigma) and viral supernatant. Lentiviral supernatant was added at concentration of 1000 MOI in a total volume of 200 ul in a 96-well U-bottom plate. The plate was spun at 2500 rpm for 60 min at room temperature. After this, cells were cultured at 37°C and 5% CO_2_ for 36 hours. Transduced cells (viable GFP^+^) were sorted on a FACSAria (BD Biosciences). MPP4s were resuspended in IMDM (Gibco) supplemented with 10% FBS, interleukin-3 (IL-3, 10 ng/ml), interleukin-6 (IL-6, 10 ug/ml), interleukin-7 (IL-7, 20 ng/ul), SCF (100 ng/ml), leukemia inhibitory factor (LIF, 20 ng/ml) and polybrene (5 ug/ml). Lentiviral supernatant was added at concentration of 100 MOI. Cells were cultured at 37°C and 5% CO_2_ for 36 hours. Transduced cells (viable GFP^+^) were sorted on a FACSAria (BD Biosciences).

### Colony Forming Unit (CFU) Assays

For B-lymphoid CFU assays, 100 GFP^+^ cells from transduced MPP4 cells were plated in Methocult M3630 (StemCell Technologies) supplemented with FMS-like tyrosine kinase like 3 ligand (FLT3L, 25 ng/ml) and SCF (50 ng/ml). For myeloid CFU assays, 100 GFP^+^ cells from transduced MPP4s or 200 GFP^+^ cells from transduced LT-HSCs were plated in Methocult GF M3434 (StemCell Technologies) and cultured at 37°C and 5% CO_2_. Scoring of colonies was done between days 7 and 10 using a Nikon Eclipse TS100 inverted microscope. CFU cloning efficiency was calculated as the sum of the myeloid and B-lymphoid colonies divided by the sum of the myeloid and B-lymphoid colonies in the NTC group. For analysis of replating potential, colonies were harvested and total viable counts were obtained, followed by plating of 1-3 x 10^4^ cells in Methocult GF M3434. Remaining cells were resuspended in RLT buffer (Qiagen). RNA was isolated by RNeasy Micro or RNeasy Mini kits (Qiagen) followed by cDNA synthesis using the SuperScript III First-Strand Synthesis System (Thermo Fisher Scientific).

### Real-Time PCR

Real-time PCR was performed using RT^2^ SYBR Green ROX qPCR Mastermix (Qiagen) on Viaa7 (Applied Biosystems) or QuantStudio 7 Flex (Applied Biosystems). Refer to Supplemental Table 5 for primer sequences used.

### Immunofluorescence Staining of LT-HSCs

Staining was performed as previously described (Florian et al., 2012). Sorted LT-HSCs were seeded on retronectin-coated glass coverslips in SFEMII supplemented with SCF (10 ng/ml), TPO (20 ng/ul), IGF2 (20 ng/ul), FGF (10 ng/ul) (BioLegend or StemCell Technologies) for at least 2 hours. Cells were fixed in 4% PFA (Affymetrix). Cells were washed with PBS, permeabilized with 0.2% Triton X-100 (Fisher Scientific) in PBS for 20 mins and blocked with 10% goat serum (Life Technologies) for 20 mins. Cells were stained by either α-KAT6B (NBP1-92036, Novus Bio) or α-H3K23ac (ab61234, Abcam) for one hour at room temperature. For secondary antibody, cells were stained with α-Rabbit conjugated with Alexa-568 (A-11036, Invitrogen) for one hour. Coverslips were mounted on slides with Gold Antifade with DAPI (Invitrogen). Imaging was performed with Leica SP8 confocal microscope. Z-stack images were summed and quantification of individual fluorescence intensities was performed by Fiji software (Schindelin et al., 2012). Scale bars in images represent 5 um.

### In Vivo Transplantation

200-350 transduced GFP^+^ cells were combined with 5 x 10^5^ bone marrow mononuclear cells (MNCs) from B6.CD45.1 mice and retro-orbitally injected into recipient B6.CD45.1 mice after 1000 rads gamma irradiation (split dose). Peripheral blood from recipient mice was analyzed by flow cytometry every 4 weeks after transplantation. Blood was collected by retro-orbital bleeding and stained with a combination of fluorochrome-conjugated antibodies from BioLegend or BD Horizon with or without red blood lysis: CD45.1 (clone A201.7), CD45.2 (clone 104), B220 (clone RA3-6B2), CD3e (clone 145-2C11), CD11b (clone M1170), Ly6g (clone 1A8), Ly6c (clone HK1.4), Ter-119 (clone TER-119), GR1 (clone RB6-8C5), CD4 (clone RM4-5), CD8a (clone 53-6.72) and CD41 (clone MWReg30). Stained cells were run on a FACSymphony A5 (BD Biosciences) and data was analyzed using FlowJo software (FlowJo, LLC).

### RNA-Seq

Transduced GFP^+^ LT-HSCs from 3 independent biological replicates were sorted directly into 350 ul of RLT buffer (Qiagen). Total RNA was isolated from cells using the RNeasy Micro kit (Qiagen), according to manufacturers’ protocols, including the optional DNase digest step. Sample concentration and quality were assessed using the Nanodrop 2000 spectrophotometer (Thermo Scientific) and the RNA 6000 Pico LabChip assay (Agilent Technologies). Libraries were prepared by the Genome Technologies core facility at The Jackson Laboratory using the Ovation RNA-seq System V2 (Nugen) and Hyper Prep Kit (Kapa Biosystems). Briefly, the protocol entails first and second strand cDNA synthesis and cDNA amplification utilizing Nugen’s Ribo-SPIA technology, cDNA fragmentation, ligation of Illumina-specific adapters containing a unique barcode sequence for each library, and PCR amplification. Libraries were checked for quality and concentration using the D5000 ScreenTape assay (Agilent Technologies) and quantitative PCR (Kapa Biosystems), according to the manufacturers’ instructions. Libraries were pooled and sequenced 75 bp single-end on the NextSeq 500 (Illumina) using NextSeq High Output Kit v2.5 reagents (Illumina). Raw data is in process of being deposited into Gene Expression Omnibus (GEO accession #TBD).

### RNA-Seq Analysis

Trimmed alignment files (with trimmed base quality value < 30, and 70% of read bases surpassing that threshold) were processed using the RSEM (v1.2.12; RNA-Seq by Expectation-Maximization) software (Li and Dewey, 2011). Alignment was completed using Bowtie 2 (v2.2.0) (Langmead and Salzberg, 2012) and processed using SAMtools (v0.1.18) (Li et al., 2009). Fragment length mean was set to 280 and standard deviation to 50. Expected read counts per gene produced by RSEM were rounded to integer values, filtered to include only genes that have at least two samples within a sample group having a cpm > 1.0, and were passed to edgeR (v3.5.3) (Robinson et al., 2010) for differential expression analysis. The Cox-Reid profile-adjusted likelihood method was used to derive tagwise dispersion estimates. The GLM likelihood ratio test was used for differential expression in pairwise comparisons between sample groups which produced exact p-values per test. The Benjamini and Hochberg’s algorithm (p-value adjustment) was used to control the false discovery rate (FDR). Features with an FDR-adjusted p-value < 0.05 were declared significantly differentially expressed.

## Acknowledgments

This work was supported by the National Institutes of Diabetes and Digestive and Kidney Diseases (NIDDK) grants R01DK118072 and R56DK112947, the Ellison Medical Foundation New Scholar Award in Aging and pilot funds from The Jackson Laboratory’s Nathan Shock Center for Excellence in the Basic Biology of Aging (P30AG038070) (J.J.T.). E.S.K. was supported by Thurgood Marshall Tuition Scholarship from University of Maine. K.Y. received support from T32HD007065, the Pyewacket Fund and the American Society of Hematology (ASH) Scholar Award. We thank Tina Mujica, Jennifer SanMiguel, Olivia Erickson, Tara Murphy and Kaiden Waldron-Francis for technical help, experimental and laboratory support and members of Trowbridge Lab for helpful discussions and critical comments. We acknowledge Rick Maser and the Genetic Engineering Technology Core, Philipp Heinrich and the Microscopy Core, Heidi Munger and the Genome Technologies Core, and Will Schott and the Flow Cytometry Core for their contribution to these studies.

