## Supplemental Materials for "Aging-Associated Decrease in the Histone Acetyltransferase KAT6B Causes Myeloid-Biased Hematopoietic Stem Cell Differentiation"

**Supplemental Figure 1. Validation of shRNA constructs used for screening epigenetic regulators of myeloid versus B-lymphoid differentiation.** (A) GO enrichment analysis of 2,766 genes identified as differentially expressed between GMP versus CLP. (B) Relative expression of shRNA target genes following knockdown in NIH/3T3 cells. Bars represent mean of $n = 3$ technical replicates.

**Supplemental Figure 2. Expression of *Kat6b* in hematopoietic stem and progenitor cell aging.** (A) Relative expression of *Kat6b* in MPP4 cells isolated from young (2-4 month) and aged (20-23 month) mice. Bars represent mean ± SEM of $n = 3$ biological replicates. (B) Log2FC of *KAT6B* expression in aged versus young human HSCs (data from Adelman et al., 2019).

**Supplemental Figure 3. Multilineage engraftment of LT-HSCs following *Kat6b* knockdown.** (A) Frequency of B ($B220^+$) and T ($CD3e^+$) cells within donor-derived PB cells ($CD45.2^+$ $GFP^+$) at one month post-transplant. Each dot represents one recipient mouse. Lines represent mean ± SEM of $n \geq 7$ biological replicates. *P* values calculated by unpaired *t* test. (B) Frequency of myeloid ($CD11b^+$), erythroid ($Ter119^+$), B ($B220^+$) and T ($CD3e^+$) cells within donor-derived PB cells at four months post-transplant. Each dot represents one recipient mouse. Lines represent mean ± SEM of $n = 3$ biological replicates. *P* values calculated by unpaired *t* test.

**Supplemental Figure 4. *Kat6b* knockdown alters gene expression programs critical for multilineage differentiation.** Top gene ontology (GO) terms enriched in genes found to be significantly downregulated in *Kat6b* sh1 versus NTC (fold change > 2 and $P < 0.05$).

**Supplemental Figure 5: Alterations in H3K23ac with aging.** (A) Representative immunofluorescence images of H3K23ac and DAPI in LT-HSCs isolated from young and aged mice. Scale bar equals 5 um. (B) Violin plots of mean fluorescence intensity (MFI) of H3K23ac in LT-HSCs isolated from young and aged mice. Solid lines indicate median and dotted lines indicate quartiles. Data points include $n$ = 31-70 individual cells sampled from $n$ = 4 biological replicate animals. $P$ values calculated by unpaired $t$ test.

**Supplemental Table 1. Sixteen Chromatin Regulatory Genes for shRNA Screening Identified Using Unbiased Differential Expression Approach**

| **Gene Symbol** | **Gene Name** |
|---|---|
| *Kat6b* | K(lysine) acetyltransferase 6B |
| *Kmt5a* | lysine methyltransferase 5A |
| *Tbl1x* | transducin (beta)-like 1 X-linked |
| *Kdm5b* | lysine (K)-specific demethylase 5B |
| *Ncor2* | nuclear receptor co-repressor 2 |
| *Suv39h2* | suppressor of variegation 3-9 2 |
| *Mta3* | metastasis associated 3 |
| *Carm1* | coactivator-associated arginine methyltransferase 1 |
| *Dot1l* | DOT1-like, histone H3 methyltransferase (S. cerevisiae) |
| *Kmt2e* | lysine (K)-specific methyltransferase 2E |
| *Arid1a* | AT rich interactive domain 1A (SWI-like) |
| *Kdm5c* | lysine(K)-specific demethylase 5C |
| *Smarcc2* | SWI/SNF related, matrix associated, actin dependent regulator of chromatin, subfamily c, member 2 |
| *Sap30l* | SAP30-like |
| *Jmjd6* | jumonji domain containing 6 |
| *Padi2* | peptidyl arginine deiminase, type II |

**Supplemental Table 2. Eight Chromatin Regulatory Genes for shRNA Screening Identified Using Candidate Gene Approach**

| **Gene Symbol** | **Gene Name** |
|---|---|
| *Rnf40* | ring finger protein 40 |
| *Atxn7l3* | ataxin 7-like 3 |
| *Prdm16* | PR domain containing 16 |

| | |
|---|---|
| *Cxxc1* | CXXC finger 1 (PHD domain) |
| *Dach1* | dashshund family transcription factor 1 |
| *Atxn7l1* | ataxin 7-like 1 |
| *Ndn* | necdin |
| *Ezh1* | enhancer of zeste 1 polycomb repressive complex 2 subunit |

**Supplemental Table 3. Gene Sets Utilized for Gene Set Enrichment Analysis (GSEA) of NTC vs. *Kat6b* shRNA RNA-Seq Data.** (see Excel file)

**Supplemental Table 4. shRNA Plasmids**

| Target Gene Symbol | Target Gene Name | Clone Number (Sigma) |
|---|---|---|
| NTC | MISSION® TRC2 pLKO.5-puro Non-Mammalian shRNA Control Plasmid DNA | SHC202 |
| *Kat6b* sh1 | K(lysine) acetyltransferase 6B | TRCN0000287544 |
| *Kat6b* sh2 | K(lysine) acetyltransferase 6B | TRCN0000039341 |
| *Crebbp* | CREB binding protein | TRCN0000231201 |
| *Rnf40* | ring finger protein 40 | TRCN0000041048 |
| *Kmt5a* | lysine methyltransferase 5A | TRCN0000241070 |
| *Atxn7l3* | ataxin 7-like 3 | TRCN0000251735 |
| *Tbl1x* | transducin (beta)-like 1 X-linked | TRCN0000109359 |
| *Atxn7l1* | ataxin 7-like 1 | TRCN0000348933 |
| *Ndn* | necdin | TRCN0000312951 |
| *Ezh1* | enhancer of zeste 1 polycomb repressive complex 2 subunit | TRCN0000313461 |
| *Kdm5b* | lysine (K)-specific demethylase 5B | TRCN0000113491 |
| *Ncor2* | nuclear receptor co-repressor 2 | TRCN0000095281 |
| *Suv39h2* | suppressor of variegation 3-9 2 | TRCN0000353741 |
| *Prdm16* | PR domain containing 16 | TRCN0000433533 |
| *Cxxc1* | CXXC finger 1 (PHD domain) | TRCN0000241393 |
| *Dach1* | dashshund family transcription factor 1 | TRCN0000075462 |

**Supplemental Table 5. Primer Sequences**

| Target | Purpose | Forward Primer (5') | Reverse Primer (5') |
|---|---|---|---|
| *B2m* | Real-time PCR | CAGTATGTTCGGCTTCCCATTC | TTCTGGTGCTTGTCTCACTGA |
| *Kat6b* set 1 | Real-time PCR | AGAAGAAAAGGGGTCGTAAACG | CAGTATGTTCGGCTTCCCATTC |
| *Kat6b* set 2 | Real-time PCR | AGCTTCTGTTTGGGGACTAAAG | GTGTCCACTACTGCCACAATC |
| eGFP | Cloning | GCAGTCGGCTCCCTCGTTGACCGA | CTGCACGCTCCCGTCCTCGATGTT |
| *Crebbp* | Real-time PCR | CCAAACGAGCCAAACTCAGC | TTTGGACGCAGCATCTGGAA |
| *Rnf40* | Real-time PCR | GACCCTACGGTGACGGAAGT | CCAGTAGCGGTTGACGATGT |
| *Kmt5a* | Real-time PCR | CAGACCAAACTGCACGACATC | CTTGCTTCGGTCCCCATAGT |
| *Atxn7l3* | Real-time PCR | AAGGAGTGTGTTTGCCCCAA | AGACTTGGATCTTCGAGGGGA |
| *Tbl1x* | Real-time PCR | CACAAGTTGCACGGCTCG | ACTGTGGCTTTACTCGGTGG |
| *Atxn7l1* | Real-time PCR | CAAGCCCTAGAACAGCGTCA | AGCAAGTTTCTGCCCTCACA |
| *Ndn* | Real-time PCR | CCAGAGGAGCTAGACAGGGT | ACGCCTGGGGATCTTTCTTG |
| *Ezh1* | Real-time PCR | CAACACTTCCCGCTGCATTC | GGCGCTTCCGTTTTCTTGTT |
| *Kdm5b* | Real-time PCR | CGAGCTGGGAAGAGTTCGC | ATCACAAGCGAATGGTGGCT |
| *Ncor2* | Real-time PCR | CCTGGTGGAAGTTCGTGGAC | ATGGTACTGGCGCTGTGTCAG |
| *Suv39h2* | Real-time PCR | GACCGCGCCAGTTTGAATG | CTAAAGGTGGGCCCTCCAAG |
| *Prdm16* | Real-time PCR | ATGGATCCCATCTACAGGGTA | CATTGCATATGCCTCCGGGT |
| *Cxxc1* | Real-time PCR | CCAAACGAGCCAAACTCAGC | TTTGGACGCAGCATCTGGAA |
| *Dach1* | Real-time PCR | GGCTTTCGACCTGTTCCTGA | AGGAAGTTCCAGTCCAACACT |

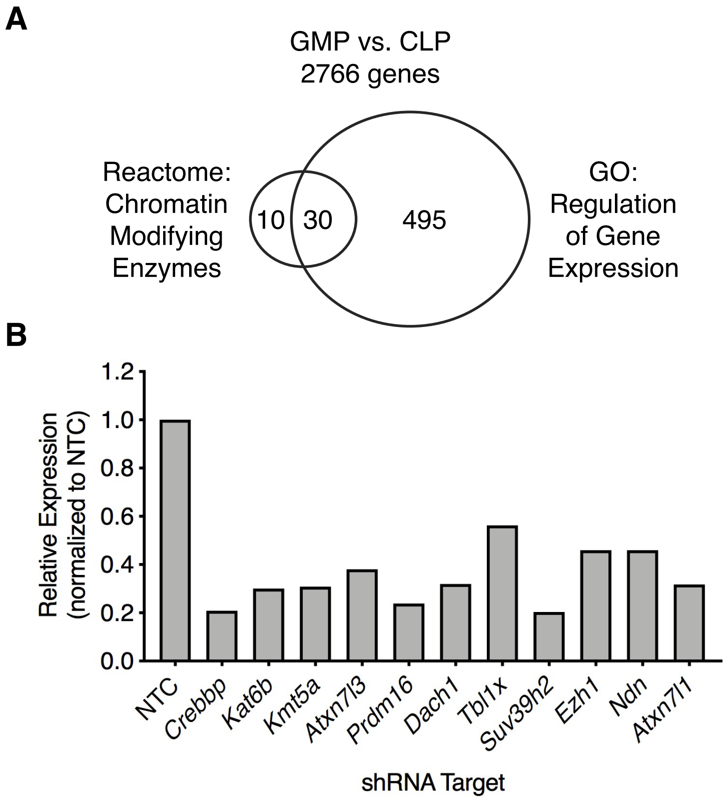
A
GMP vs. CLP
2766 genes
Reactome:
Chromatin
Modifying
Enzymes
10
30
495
GO:
Regulation
of Gene
Expression
B
Relative Expression
(normalized to NTC)
1.2
1.0
0.8
0.6
0.4
0.2
0.0
NTC
Crebbp
Kat6b
Kmt5a
Atxn7l3
Prdm16
Dach1
Tbl1x
Suv39h2
Ezh1
Ndn
Atxn7l1
shRNA Target

Supplemental Figure 2

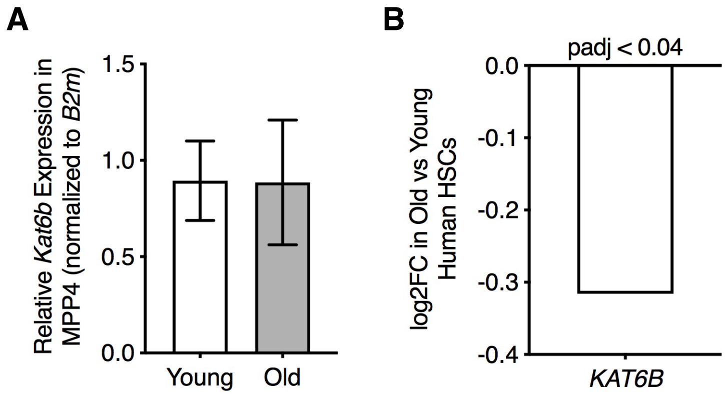

### Supplemental Figure 3

A

B

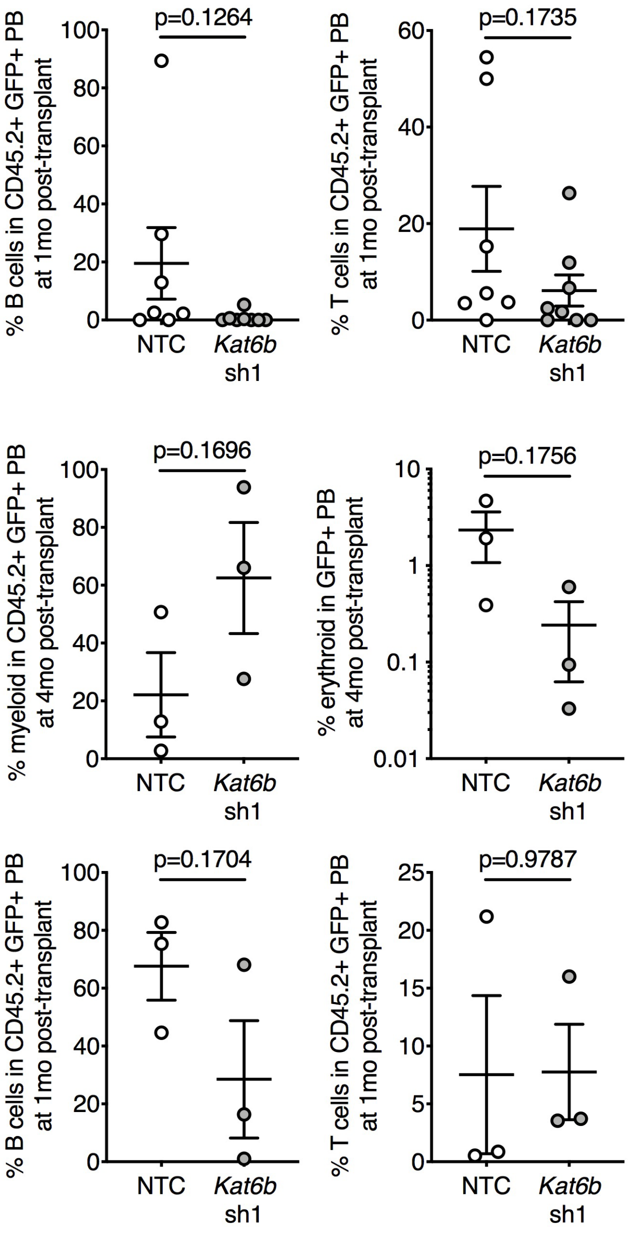

### Supplemental Figure 4

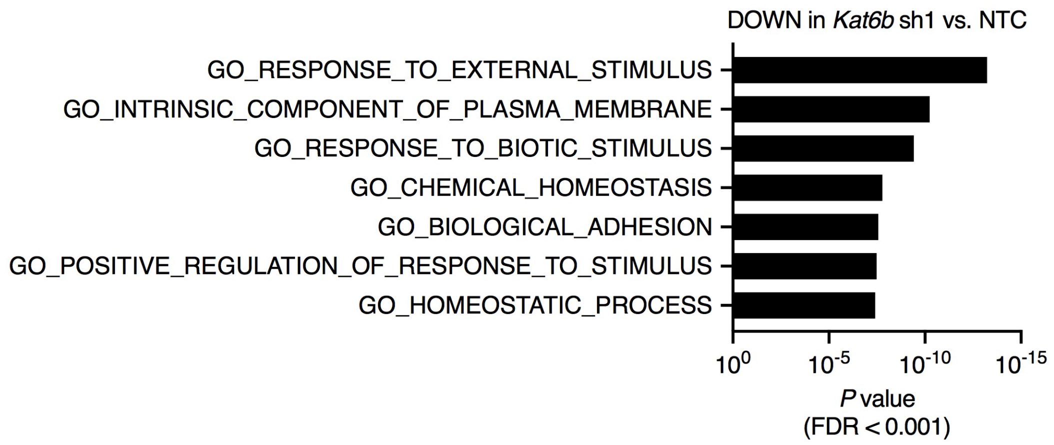

Supplemental Figure 5

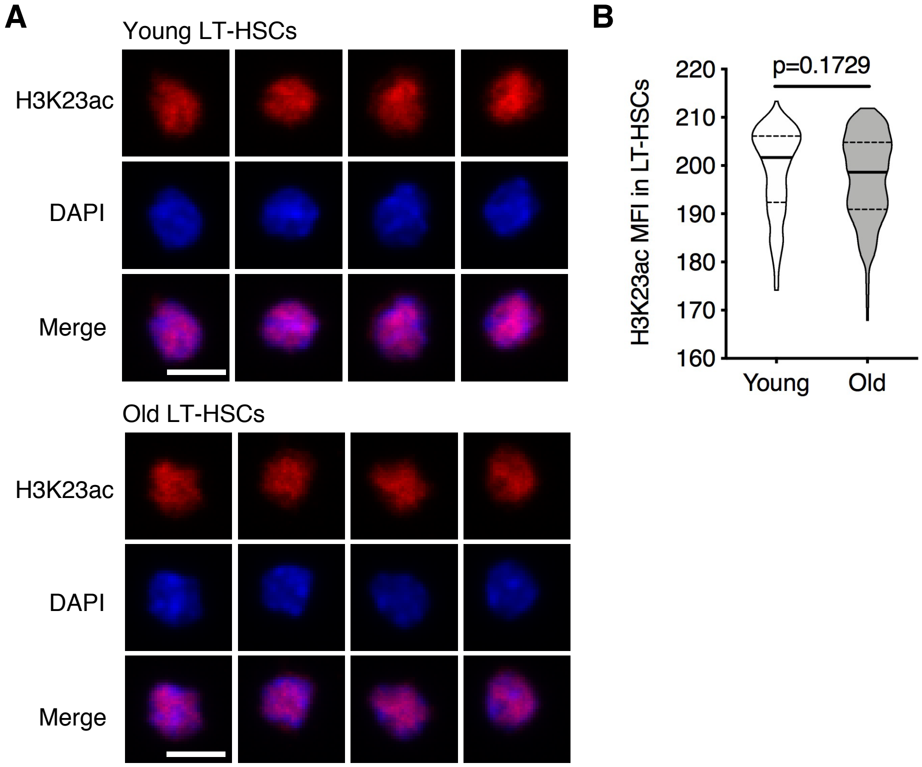
